# Prophylactic Lipoxin A_4_ Attenuates *Clostridioides difficile* Infection by Augmenting Epithelial Barrier and Resolving Inflammation

**DOI:** 10.64898/2026.02.03.703554

**Authors:** Hui Wen, Yunqing Xiang, Yue Yu, Zhixin Ma, Ying Xin, Yufang Deng, Huipai Peng, Yong Shi, Nan Li, Shuqiang Huang

**Affiliations:** State Key Laboratory of Quantitative Synthetic Biology, Shenzhen Institute of Synthetic Biology, Shenzhen Institutes of Advanced Technology, Chinese Academy of Sciences, Shenzhen, 518055, China; University of Chinese Academy of Sciences, Beijing, 100049, China; Faculty of Synthetic Biology, Shenzhen University of Advanced Technology, Shenzhen, 518107, China; Department of Mechanical, Materials and Manufacturing Engineering, University of Nottingham Ningbo China, Ningbo, 315100, China

## Abstract

*Clostridioides difficile* infection (CDI) is a leading healthcare-associated diarrhea with high recurrence rates, partially due to antibiotic-induced dysbiosis and dysregulated host inflammation. Specialized pro-resolving mediators (SPMs), such as Lipoxin A_4_ (LXA_4_), offer promise in controlling excessive inflammation and promoting tissue repair, yet their role in CDI remains unexplored. Here, we developed a compact, gas-tight gut-on-a-chip (GOC) system that reconciles the anaerobic requirements of *C. difficile* with the oxic needs of human intestinal epithelium, enabling physiologically relevant co-culture within a standard incubator. A CDI *in vitro* model was established based on this GOC system. Using the model, we demonstrated that prophylactic administration of LXA_4_ significantly preserved epithelial barrier integrity, attenuated pro-inflammatory cytokine secretion (IL-8 and IFN-γ), and reduced bacterial colonization. Transcriptomic analysis revealed that LXA_4_ pretreatment upregulated genes involved in cell junction organization while downregulated immune activation pathways. These protective effects were validated in a murine CDI model, where LXA_4_ pretreatment reduced weight loss, pathological damage, and fecal bacterial burden. Furthermore, prophylactic administration of LXA_4_ synergized with vancomycin treatment further enhanced antibiotic efficacy while allowing a 50% dose reduction without compromising therapeutic outcomes. Our study establishes a robust approach for CDI research and highlights the prophylactic and adjuvant potential of inflammation-resolving strategies, offering a novel approach to mitigate CDI incidence and improve treatment outcomes.

## Introduction

*Clostridioides difficile* is an opportunistic intestinal pathogen and the primary cause of antibiotic-associated diarrhea ***Rupnik et al. (2009***). Under the conditions of gut microbiota dysbiosis, *C. difficile* infection (CDI) exhibits high incidence rates and mortality, establishing it as one of the most prevalent healthcare-associated infections ***Barbut et al. (2000); Guh et al. (2020***).

Oral antibiotics such as vancomycin or fidaxomicin are the standard first-line therapy for CDI ***Bainum et al. (2023***). However, this approach creates a therapeutic paradox. While reducing the pathogen load, these agents further deplete the already compromised gut microbiota and exacerbate dysbiosis ***Crowther and Wilcox (2015***). Consequently, antibiotic treatment is associated with a high recurrence rate, affecting between 20% and 30% of patients after an initial episode ***Bainum et al. (2023***). Additionally, sustained antibiotic pressure raises concerns for the development of antimicrobial resistance, presenting a significant long-term public health challenge ***Murray et al. (2022***). This compels the search for more effective and safer therapeutic strategies.

The pathogenesis of CDI critically involves a dysregulated host inflammatory response that directly exacerbates intestinal injury ***Wang et al. (2024***). *C. difficile* secreted toxins not only disrupt the epithelial barrier but also actively dysregulate host immune signaling within the intestinal tissue, triggering an excessive release of inflammatory mediators ***Pourliotopoulou et al. (2024***). This process can be amplified by pre-existing dysbiosis, which compromises innate immune surveillance ***Spigaglia (2024***). This disproportionate inflammatory cascade inflicts significant collateral damage on the mucosal architecture, compromising both tissue integrity and function ***Kordus et al. (2022***). Importantly, the resulting damaged intestinal environment creates a pathophysiological milieu that further facilitates *C. difficile* colonization and toxin production ***Wang et al. (2024***). Thus, a self-perpetuating cycle is established within the host: toxin-driven inflammation causes tissue damage, and the damage in turn promotes further bacterial proliferation and pathological progression. Consequently, we propose that a therapeutic strategy aimed at controlling excessive inflammation and re-establishing a balanced immune response may offer a breakthrough for more effective CDI management.

Acute inflammation is typically self-limiting, followed by an active resolution phase mediated by specialized pro-resolving mediators (SPMs) that restore tissue homeostasis ***Serhan (2014); Fuller-ton and Gilroy (2016); Serhan and Petasis (2011***). Lipoxin A_4_ (LXA_4_) is one of the most well-characterized SPMs, derived from arachidonic acid via transcellular biosynthesis. It potently inhibits neutrophil chemotaxis and infiltration, stimulates non-phlogistic phagocytosis of apoptotic cells by macrophages, and promotes mucosal repair and barrier function ***Gronert et al. (1998); Gewirtz et al. (2002***). These properties suggest that LXA_4_ could be a promising therapeutic agent for suppressing excessive acute inflammatory responses in CDI. Previous studies have shown that LXA_4_ analogs can selectively inhibit Salmonella typhimurium–induced secretion of neutrophil-attracting chemokines in intestinal epithelial cells, thereby dampening the inflammatory cascade ***Gewirtz et al. (1998***). Additionally, LXA_4_ was found to enhance bronchial epithelial barrier function in cystic fibrosis by promoting tight junction assembly during *Pseudomonas aeruginosa* infection ***Higgins et al. (2016***). Based on these evidences, we hypothesize that LXA_4_ may alleviate CDI pathology by modulating host immune responses. Therefore, targeted use of LXA_4_ could offer a novel interventional strategy for CDI. However, the specific effects of LXA_4_ on CDI remain unexplored.

The pathophysiology of CDI has been extensively investigated using murine models ***Fachi et al. (2024***). However, interspecies differences can significantly influence immune responses, host-pathogen interactions, and the pharmacokinetics and efficacy of therapeutic agents ***Calvigioni et al. (2023***). This limits the direct translatability of findings to humans and has motivated the development of advanced in vitro human intestinal models. These models hold promise for more accurately recapitulating the specific host-microbe interactions within the human gut ***Ewin et al. (2023***). Conventional in vitro models for CDI primarily utilize static monolayer cultures of immortalized epithelial cells, grown in multi-well plates or Transwell inserts ***Jafari et al. (2016); Janvilisri et al. (2010***). A fundamental limitation of this approach is the conflicting environmental requirements, as *C. difficile* thrives under strict anaerobic conditions while epithelial cells require normoxic conditions for optimal growth. Consequently, such setups are restricted to short-term co-culture experiments following bacterial inoculation. This brief exposure window is insufficient for studying the progressive cellular and molecular events that characterize CDI pathophysiology. Alternatively, many existing models rely on the static application of purified *C. difficile* toxins rather than live bacteria ***Kasendra et al. (2014); Leslie et al. (2015***). While informative for toxin biology, this method fails to capture the dynamic and multi-faceted interactions between live bacteria and the host epithelium. These interactions are critical for a comprehensive understanding of infection dynamics.

To reconcile the divergent oxygen demands, specialized devices featuring two chambers separated by a porous membrane have been developed ***Bang et al. (2025); Anonye et al. (2019***). In these systems, an oxic chamber supports epithelial cells and an adjacent anoxic chamber cultivates *C. difficile*. Furthermore, such configurations can provide epithelial cells with fluidic shear stress, a key mechanical cue that promotes phenotypic differentiation and maturation. Typically constructed from rigid plastics, these devices remain compatible with standard cell culture incubators. Also based on the two chambers oxygen segregation principle, gut-on-a-chip (GOC) models fabricated from elastic silicone elastomers have emerged as superior platforms for mimicking the native intestinal microenvironment ***Jalili-Firoozinezhad et al. (2019); Meza-Torres et al. (2025***). The elastic property enables GOC to integrate more essential physiological cues, such as cyclic peristalsis-like mechanical strain. This makes GOC systems uniquely drive intestinal epithelial development toward a more physiologically relevant state, closely mirroring the structural complexity and functional maturity observed *in vivo*. Nonetheless, a significant practical challenge remains due to the high gas permeability of the silicone elastomer. To maintain the stringent anoxia required for anaerobic bacteria, the entire GOC device must be housed within a custom-built anaerobic chamber. This requirement introduces practical challenges related to cost, complexity, and compatibility with routine laboratory workflows.

Herein, we developed a GOC-based system to establish an in vitro CDI model. The system is configured within a sealed chamber that can be placed in a standard cell culture incubator, offering a compact, cost-effective, and operationally straightforward approach for co-culture experiments with reduced risk of microbial dissemination. Employing this system, we investigated the role of the pro-resolving mediator LXA_4_ in CDI. We found that while LXA_4_ administration after infection did not show direct therapeutic efficacy, its prior application significantly alleviated the severity of CDI pathology, represented as LXA_4_ treatment greatly strengthened the epithelial barrier and mitigated dysregulated inflammatory responses. This protective effect was corroborated in a murine model. Furthermore, we revealed a promising synergistic strategy wherein the combination of LXA_4_ with standard antibiotics enhanced therapeutic efficacy beyond antibiotic monotherapy. Collectively, our work not only provides a robust and accessible platform for studying host-C. difficile interactions but also opens new avenues for developing inflammation resolution-based prophylactic and adjunctive strategies to augment current treatments for healthcare-associated infections.

## Results

### Establishment of a CDI model on a GOC system

A key advancement of our platform is its compact and fully integrated design, which reconciles the conflicting oxygen requirements for epithelial cells and *C. difficile* within a single, sealable chamber compatible with standard incubators. The setup of the GOC-based system is shown in ***Figure 1***A. It consists of four main components: an anaerobic gas-generating sachet to establish an anoxic atmosphere, an oxygen indicator to monitor oxygen levels, a GOC microfluidic chip that mimics the human intestinal tract, and a gas-tight transparent chamber that isolates all components from the ambient normoxic environment. Culture media with different oxygen partial pressures are maintained outside the chamber and perfused into the GOC device through two sealed ports on its sidewall. Effluent media are collected separately inside the chamber to facilitate subsequent recovery and analysis. The entire assembly is compact, occupying an area of < 300 cm^2^, and can be readily placed inside a conventional cell culture incubator. The gas-tight design prevents bacterial escape, allowing safe operation alongside other cell cultures in the same workspace.

**Figure 1.**
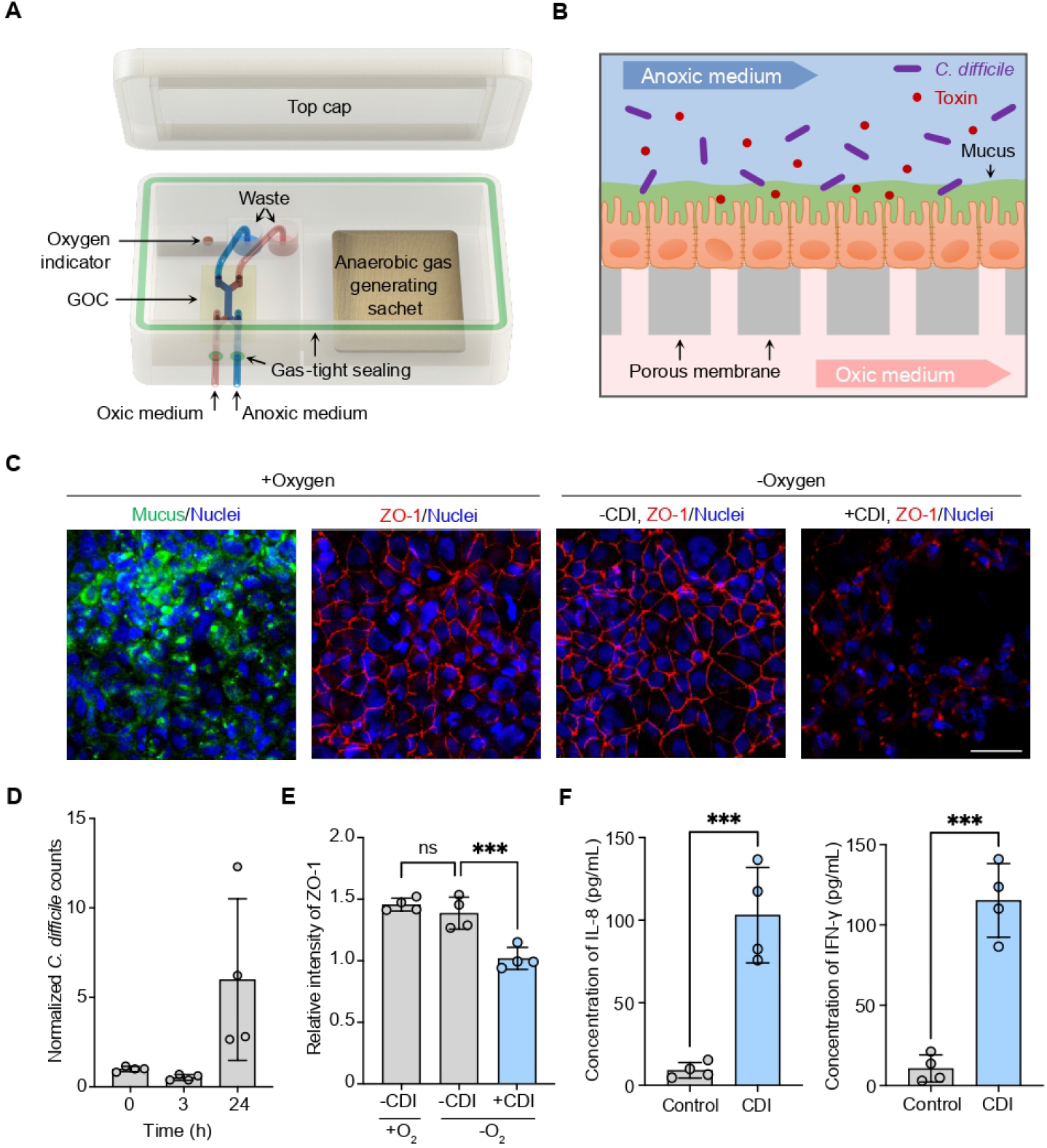
Modeling CDI using a GOC-based system. (A) Schematic of the GOC system design. (B) Schematic of the GOC-based *in vitro* CDI model. (C) Confocal micrographs of the intestinal epithelium cultured for 24 hours in the GOC. The upper microchannel was perfused with normoxic (+ Oxygen) or anaerobic (-Oxygen) culture medium. Under anaerobic condition, the intestinal epithelium was either infected (+ CDI) or not infected (– CDI) by *C. difficile*. Tight junctions were immunostained for ZO-1 (red), the mucus layer was labeled with wheat germ agglutinin (WGA, green), and nuclei were stained with DAPI (blue). Scale bar: 50 µm. (D-F) Quantitative analysis of (D) *C. difficile* cell counts in the GOCs at 0, 3 and 24 hours post-infection; (E) the average ZO-1 fluorescence intensity of intestinal epithelium in GOCs; and (F) secretion of pro-inflammatory cytokines IL-8 and IFN-γ in GOCs with or without *C. difficile* infection. Data are presented as mean ± SD (n = 4 independent replicates). Significance was determined by unpaired t-test (* P < 0.05, ** P < 0.01, *** P < 0.001). **Figure 1—figure supplement 1**. Characterization of the GOC device and on-chip cell culture.

We employed a widely adopted GOC design to establish the CDI model ***Huh et al. (2013***). As illustrated in ***Figure 1***B, the GOC chip comprises two vertically stacked microchannels separated by a porous membrane (***Figure 1—figure Supplement 1***A). Caco-2 cells, derived from human colorectal adenocarcinoma, were seeded onto the membrane and cultured to form a confluent monolayer resembling the intestinal epithelium (***Figure 1—figure Supplement 1***B). This cell layer physically separates the two microchannels, permitting the maintenance of an anoxic environment in the upper channel and an oxic environment in the lower channel. The pore size and density of the membrane allow the free diffusion of oxygenated mammalian cell culture medium to the basal side of the Caco-2 monolayer while preventing cell migration into the lower channel. *C. difficile* suspended in anoxic medium was introduced into the upper channel to enable bacterial colonization and proliferation, thereby simulating infection. Culture medium was continuously perfused at a rate of 30 µL/h to mimic intestinal fluid flow and shear stress.

The formation of intercellular tight junctions and the secretion of mucus are critical indicators of functional intestinal epithelium in vitro. To evaluate our GOC system, we characterized the tight junction protein Zonula Occludens-1 (ZO-1) and mucin production after 72 hours of oxic Caco-2 culture. Mucin secretion confirmed Caco-2 cell differentiation, and robust ZO-1 expression indicated the establishment of a functional epithelial barrier (***Figure 1***C). The oxic medium was then replaced with anoxic medium to simulate the anaerobic luminal environment of the human intestine. After 24 hours under anoxic conditions, no adverse effects on epithelial cells were observed, as evidenced by comparable ZO-1 intensity (***Figure 1***C, E) and cell viability (***Figure 1—figure Supplement 1***C) between anoxic and normoxic cultures.

To establish the CDI model, *C. difficile* was inoculated into the upper channel of the GOC device. Following 24 hours of coculture, bacterial counts increased near 6-fold (***Figure 1***D), confirming the suitability of the anoxic environment for *C. difficile* growth. Importantly, infection significantly disrupted epithelial tight junctions, as shown by markedly reduced ZO-1 signal (***Figure 1***C, E), indicating compromised barrier integrity. We further assessed the inflammatory response triggered by *C. difficile*. Levels of the representative pro-inflammatory cytokines IL-8 and IFN-γ increased approximately 14-fold and 12-fold, respectively, compared to the non-infection state (***Figure 1***F). These results demonstrate that our GOC system successfully recapitulates key pathophysiological features of CDI, confirming the establishment of a robust in vitro CDI model.

### Prophylactic effect of LXA_4_ against CDI in in vitro model

Building on the established GOC-based in vitro CDI model, we next sought to evaluate the therapeutic and prophylactic potential of the pro-resolving mediator LXA_4_ against CDI induced intestinal injury. To first determine if LXA_4_ itself exerted any adverse effects on epithelial cells, we treated intestinal epithelial cells with 200 nM LXA_4_ for 24 hours. The treatment did not affect intercellular tight junctions indicated by ZO-1 expression remained unchanged compared with controls, proven that LXA_4_ is non toxic to epithelial cells (***Figure 2—figure Supplement 1***A). This result is consistent with previous reports. We then assessed whether LXA_4_ could cure or alleviate CDI mediated damage to the intestinal epithelium. In the treatment regimen, LXA_4_ was administered 6 hours post infection, and outcomes were analyzed 24 hours after infection (***Figure 2***A). Immunofluorescence staining for the tight junction protein ZO-1 revealed severe disruption of epithelial integrity following CDI, regardless of LXA_4_ treatment (***Figure 2***B). Quantitative analysis showed no significant difference in ZO-1 expression between the CDI control and LXA_4_ treated groups (***Figure 2***C), indicating that LXA_4_ administration after infection did not confer therapeutic benefit.

**Figure 2.**
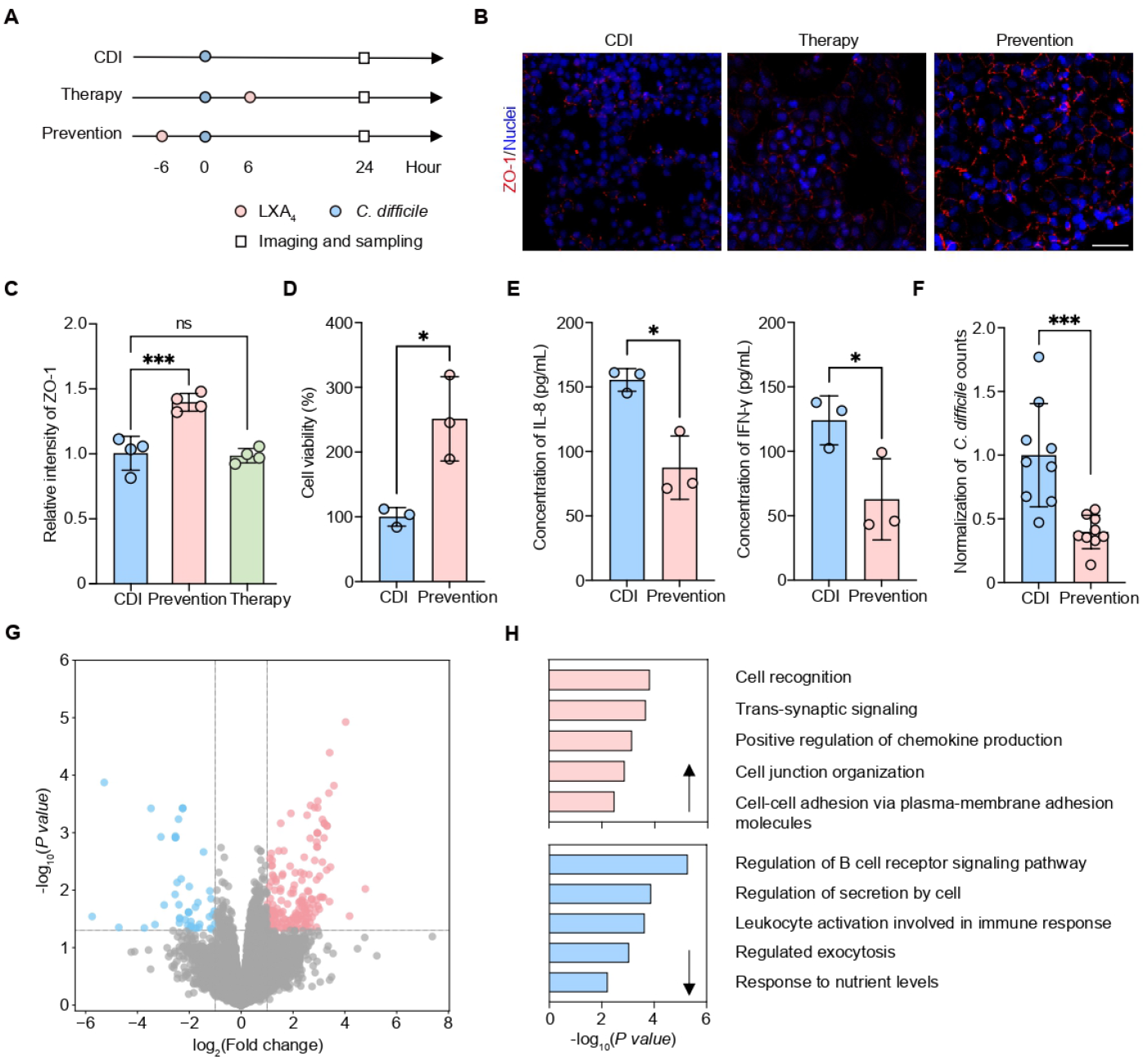
Prevention effect of LAX_4_ against CDI in the GOC-based *in vitro* system. (A) Schematic of experimental timeline for LXA_4_ administration in the control (CDI), prevention and therapy groups. (B) Confocal micrographs of the intestinal epithelium in GOCs subjected to prevention and therapy groups. Tight junctions were immunostained for ZO-1 (red) and nuclei were stained with DAPI (blue). Scale bar, 25 µm. (C-F) Quantitative analysis of (C) the average ZO-1 fluorescence intensity in the intestinal epithelium in CDI control, LXA_4_ prevention and therapy groups (n = 4 independent replicates); (D) epithelial cell viability with or without LXA_4_ prevention (n = 3 independent replicates); (E) secretion of the pro-inflammatory cytokines IL-8 and IFN-γ with or without LXA_4_ prevention (n = 3 independent replicates); and (F) *C. difficile* cell counts in GOCs with or without LXA_4_ prevention (n = 9 independent replicates). (G) Differentially expressed genes of intestinal epithelium cells from GOCs with or without LAX_4_ prevention. Genes with |log_2_(fold change)| >= 1 and adjusted P value < 0.05 were considered significantly differentially expressed. (H) GO analysis of up- and down-regulated genes in the LXA_4_ prevention group. The top five enriched terms are displayed. Data in (C–F) are presented as mean ± SD; significance was determined by unpaired t-test (* P < 0.05, ** P < 0.01, *** P < 0.001). **Figure 2—figure supplement 1**. Cytotoxicity assessment of LXA_4_ and its impact on the transcriptomic landscape.

We therefore examined the prophylactic potential of LXA_4_. In the prevention regimen, LXA_4_ was added 6 hours before bacterial challenge, with data collected 24 hours post infection (***Figure 2***A). Notably, pretreatment with LXA_4_ markedly preserved tight junction organization, as evidenced by a significantly stronger ZO-1 signal compared with both the untreated control and the post infection treatment group (***Figure 2***B, C). Epithelial cell viability was also substantially maintained; the LXA_4_ pretreated group exhibited approximately 2.5 fold higher viability than the untreated CDI control (***Figure 2***D).

Given the well established role of LXA_4_ as a pro-resolving mediator, we further asked whether it could dampen the characteristic inflammatory response triggered by CDI. As anticipated, levels of the key pro inflammatory cytokines IL 8 and IFN-γ were reduced by about 50% in the prevention group relative to the untreated control (***Figure 2***E), demonstrating a pronounced anti inflammatory effect of LXA_4_ during CDI. These data suggest a direct link between the anti inflammatory activity of LXA_4_ and its prophylactic efficacy.

It has been reported that *C. difficile* exacerbates intestinal damage by driving an excessive inflammatory response, which not only disrupts the epithelial barrier but also releases host derived nutrients and metabolites that can support bacterial proliferation ***Wang et al. (2024***). We thus hypothesize that the prophylactic effect of LXA_4_ may operate through two interrelated mechanisms: first, by attenuating the inflammatory cascade, LXA_4_ limits the availability of pro proliferative host factors for *C. difficile*; second, the preserved epithelial barrier integrity, reflected in enhanced cell viability and tight junction stability, likely hinders bacterial adhesion and colonization. Consistent with this hypothesis, bacterial counts in the LXA_4_ prevention group were approximately threefold lower than those in the untreated control (***Figure 2***F).

To gain further insight into epithelial responses to LXA_4_, we performed transcriptomic analysis comparing LXA_4_ pretreated and CDI infected epithelial layers. A total of 226 differentially expressed genes (48 downregulated, 178 upregulated) were identified in LXA_4_ exposed epithelia (***Figure 2***G, ***Figure 2—figure Supplement 1***B). Gene Ontology enrichment analysis revealed that upregulated genes were significantly associated with cell recognition, regulation of chemokine production, cell junction organization, and intercellular adhesion, indicating enhanced pathogen sensing and barrier reinforcement. Conversely, downregulated genes were enriched in immune activation pathways, consistent with a restrained inflammatory response (***Figure 2***H). These findings demonstrate that LXA_4_ reprograms the intestinal epithelial transcriptome to bolster barrier function and temper immune signaling.

Collectively, LXA_4_ can protect against CDI through prophylactic administration by preserving epithelial barrier integrity, reducing inflammation, and limiting *C. difficile* proliferation.

### Validating the prophylactic effect of LXA_4_ against CDI in an *in vivo* model

To further validate the prophylactic effect of LXA_4_ that had been initially observed under in vitro experimental conditions, we subsequently employed a murine model of CDI in order to comprehensively evaluate its protective efficacy within a living organism (***Figure 3***A). In mice that were infected with *C. difficile* but did not receive any pretreatment, the infection resulted in a pronounced and progressive decline in overall health, marked by a substantial body weight loss exceeding 5%. By contrast, those mice that had been prophylactically administered LXA_4_ exhibited a significant preservation of physiological condition, maintaining body weights that were closely comparable to those of the uninfected control group throughout the observation period, and demonstrating no measurable reduction attributable to the infection (***Figure 3***B). Furthermore, quantitative micro-biological analysis revealed that pretreatment with LXA_4_ led to a remarkable reduction in fecal *C. difficile* bacterial loads, achieving an approximate 10-fold decrease compared to the infected but untreated control animals (***Figure 3***C).

**Figure 3.**
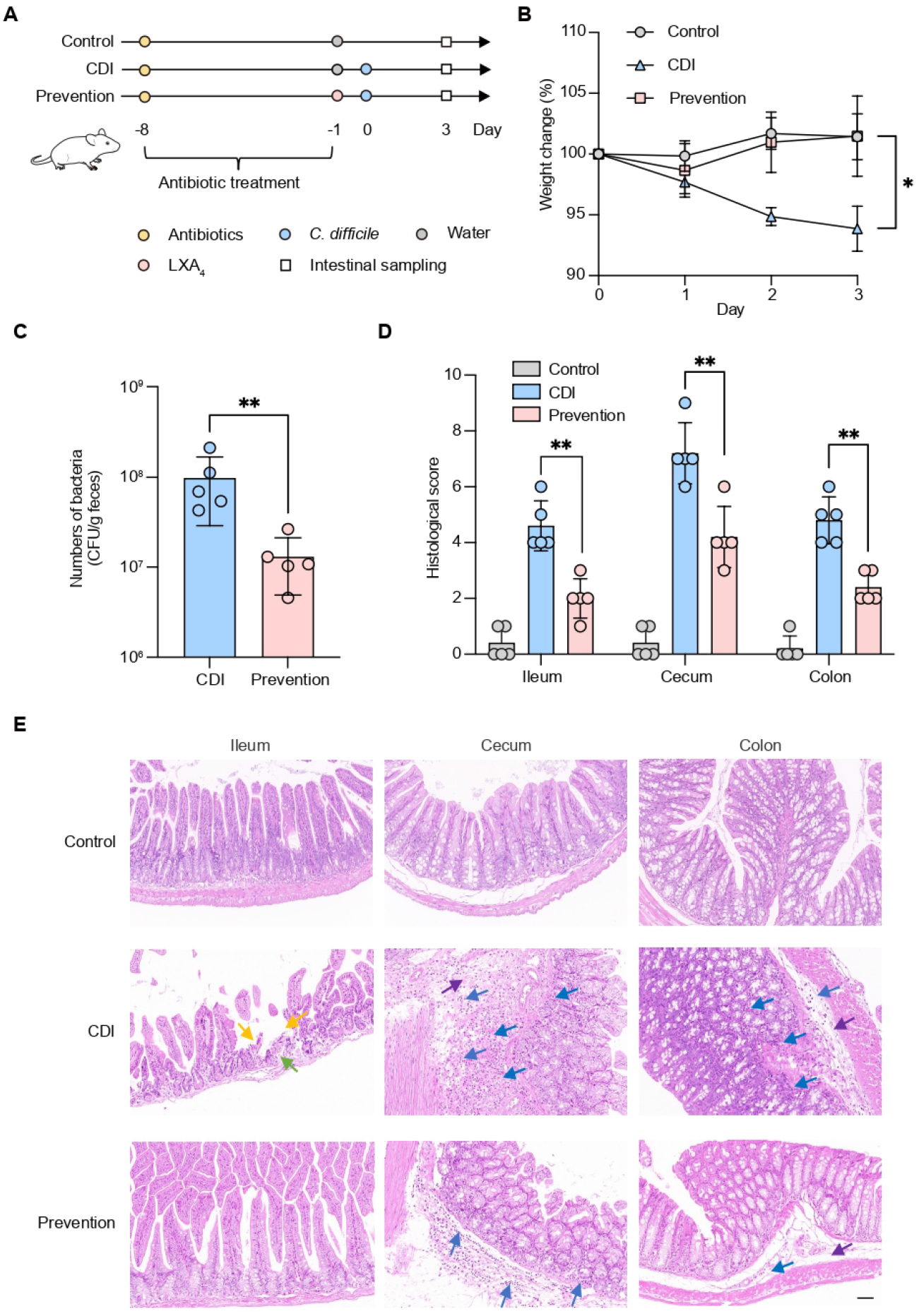
Protective effect of LXA_4_ against CDI in a murine model. (A) Experimental scheme for establishing the CDI model in mice, including the timeline of LXA_4_ administration. (B) Body weight changes in the CDI control and LXA_4_ prevention groups. (C) *C. difficile* bacterial loads in feces at day 3 post-infection with or without LXA_4_ prevention. (D) Histopathological scores of ileum, cecum, and colon from H&E-stained sections with or without LXA_4_ prevention. (E) Representative H&E-stained images of ileum, cecum, and colon tissues with or without LXA_4_ prevention. Arrows indicate key pathological features: immune cell infiltration (blue), villous epithelium loss (yellow), edema (purple), and crypt loss (green). Scale bar: 50 µm. Data in (B–D) are presented as mean ± SD (n = 5 mice per group). Statistical significance was determined by unpaired t-test (* P < 0.05, ** P < 0.01).

Histopathological examination results showed that the intestinal tissues of mice infected with *C. difficile* exhibited obvious pathological changes, mainly manifested as severe submucosal edema, massive neutrophil infiltration, and significant epithelial structural damage, including the loss of intestinal crypts and disruption of mucosal tissue architecture (***Figure 3***D, E). In contrast, the experimental group of mice that received preventive administration of LXA_4_ showed significantly improved clinical disease scores, indicating that LXA_4_ has an obvious protective effect against infection-induced pathological damage. In mice pretreated with LXA_4_, the degree of pathological changes in intestinal tissues was significantly decreased, with both the extent of epithelial damage and the number of inflammatory cell infiltrations reduced by approximately 50% to 60% (***Figure 3***D, E).

Together, these in vivo results corroborate our in vitro findings, further validating the prophylactic efficacy of LXA_4_ against CDI and its ability to alleviate associated clinical symptoms.

### Synergistic efficacy of LXA_4_ combined with antibiotics against CDI

While LXA_4_ monotherapy demonstrated limited therapeutic efficacy when administered after infection, it exhibited pronounced prophylactic potential against CDI-induced pathology. Given that antibiotics remain the first-line clinical treatment for CDI, we hypothesized that a combined strategy, which leveraging the preventive capacity of LXA_4_ alongside antibiotic therapy, could offer a more effective approach to managing healthcare-associated CDI. To test this, we selected vancomycin, a cornerstone antibiotic routinely employed in the clinical management of CDI.

To establish the baseline therapeutic response, we first evaluated vancomycin monotherapy in our GOC-based CDI model (***Figure 4***A). Vancomycin alone effectively ameliorated CDI-related damage, consistent with its established clinical activity. Compared to the infected control, vancomycin monotherapy significantly increased tight-junction protein ZO-1 expression (***Figure 4***B, C) and reduced levels of the pro-inflammatory cytokines IL-8 and IFN-γ (***Figure 4***D), confirming that our *in vitro* model reliably recapitulates the antibiotic response observed *in vivo*.

**Figure 4.**
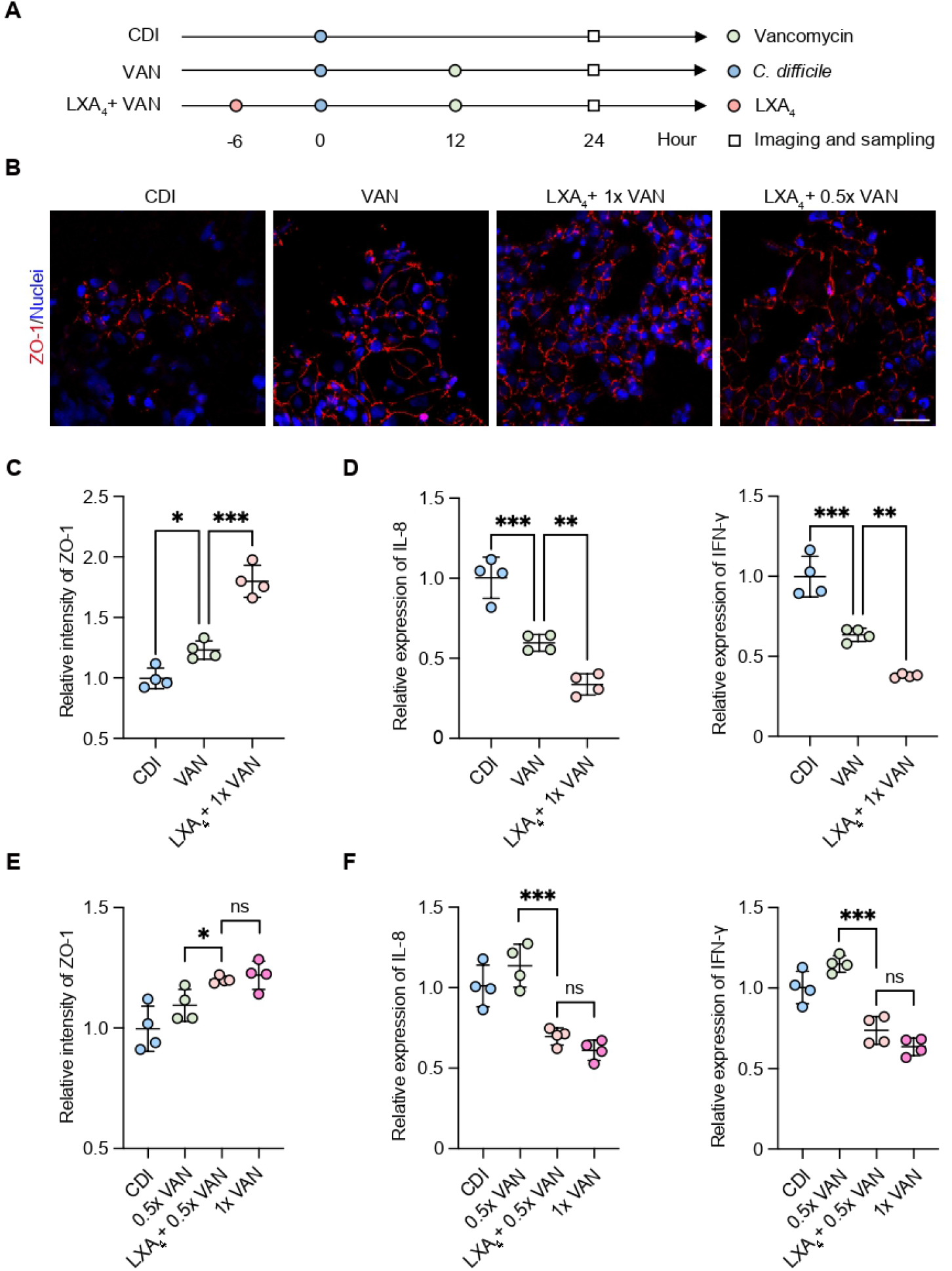
Adjunctive effect of LXA_4_ with vancomycin for the treatment of CDI. (A) Experimental scheme for the combined administration of LXA_4_ and vancomycin. (B) Confocal micrographs of the intestinal epithelium in GOCs under different conditions: CDI control, vancomycin monotherapy (VAN), LXA_4_ combined with full-dose vancomycin (LXA_4_ + 1× VAN), and LXA_4_ combined with half-dose vancomycin (LXA_4_ + 0.5× VAN). Tight junctions are immunostained for ZO-1 (red) and nuclei are stained with DAPI (blue). Scale bar: 50 µm. (C-D) Quantitative analysis of (C) the average ZO-1 fluorescence intensity and (D) the secretion of pro-inflammatory cytokines (IL-8 and IFN-γ) in GOCs from the CDI control, VAN, and LXA_4_ + 1× VAN groups. (E-F) Quantitative analysis of (E) the average ZO-1 fluorescence intensity and (F) the secretion of IL-8 and IFN-γ in GOCs from the CDI control, VAN, LXA_4_ + 1× VAN, and LXA_4_ + 0.5× VAN groups. Data in (C-F) are presented as mean ± SD (n = 4 independent replicates). Statistical significance was determined by unpaired t-test (* P < 0.05, ** P < 0.01, *** P < 0.001).

Building on this validated model, we next asked whether LXA_4_ pretreatment could further enhance therapeutic efficacy of vancomycin. In the combined regimen, LXA_4_ was administered 6 hours before *C. difficile* inoculation and vancomycin was added 12 hours post-infection, with readouts taken 24 hours after bacterial challenge (***Figure 4***A). The combination markedly improved outcomes relative to vancomycin alone. Specifically, the ZO-1 expression increased 1.4-fold (***Figure 4***B, C), while levels of the pro-inflammatory cytokines IL-8 and IFN-γ fell by approximately 3-fold and 2.6-fold (***Figure 4***D). These results demonstrated that LXA_4_ pretreatment synergized with vancomycin to strengthen epithelial barrier integrity and more effectively suppress inflammation during CDI. Thus, prophylactic LXA_4_ indeed enhanced the therapeutic effect of vancomycin against CDI *in vitro*.

Clinically, severe CDI requires higher antibiotic dosages, which increases the risk of patient toxicity and potential organ damage ***Johnson et al. (2021); McDonald et al. (2018***). Therefore, therapeutic strategies that enhance antibiotic efficacy without raising dosage are highly desirable. We investigated whether a reduced antibiotic dosage combined with prophylactic LXA_4_ could achieve efficacy comparable to that of a full-dose regimen. Experimentally, the vancomycin concentration was halved (from 0.5 to 0.25 µg/mL) in the presence of LXA_4_ pretreatment, the combination regimen achieved therapeutic outcomes comparable to those of full-dose vancomycin alone, with no statistically significant differences in ZO-1 expression or pro-inflammatory cytokine levels (***Figure 4***E, F).

Taken together, prophylactic LXA_4_ enhanced the therapeutic profile of vancomycin against CDI, enabling a substantial reduction in antibiotic dosage without compromising efficacy.

## Discussion

In this study, we developed a compact, integrated gut-on-a-chip platform that reconciles the divergent oxygen requirements of aerobic intestinal epithelium and anaerobic *C. difficile* within a single gas-tight chamber compatible with standard cell culture incubators. This design not only prevents microbial dissemination but also offers a scalable and user-friendly system for host–pathogen coculture. Using this platform, we established a robust *in vitro* model of CDI that successfully recapitulates key pathophysiological features of the infection, including disruption of epithelial tight junctions, elevated secretion of pro-inflammatory cytokines, and progressive bacterial colonization, thus providing a physiologically relevant tool for mechanistic investigation.

Our findings demonstrated that the specialized pro-resolving mediator LXA_4_ exerted significant protective effects against CDI when administrated prophylactically. LXA_4_ pretreatment strengthened epithelial barrier integrity, as evidenced by preserved ZO-1 expression and enhanced cell viability, while simultaneously dampening the release of pro-inflammatory cytokines such as IL-8 and IFN-γ. These dual actions, barrier protection and inflammation resolution, collectively limit *C. difficile* colonization and mitigate infection induced tissue damage. Notably, LXA_4_ synergized with the frontline antibiotic vancomycin, improving epithelial repair and inflammatory suppression beyond antibiotic monotherapy. Moreover, LXA_4_ pretreatment allowed a substantial reduction in vancomycin dose without compromising therapeutic efficacy, highlighting its potential as an adjuvant to decrease antibiotic exposure and associated risks.

From a clinical perspective, prophylactic LXA_4_ administration could address a critical unmet need in hospitalized patients at high risk for CDI as well, particularly those receiving broad-spectrum antibiotics, for whom no approved preventive strategies are currently available. Although several candidate vaccines targeting high-risk populations have entered clinical development, most advanced candidates (e.g., toxoid-based vaccines) require multiple-dose regimens (typically three doses) and, as demonstrated in recent phase 3 trials, have thus far shown limited success in preventing primary CDI ***Donskey et al. (2024); Kuijper and Gerding (2024***). In this context, a readily administrable prophylactic agent such as LXA_4_, or its stable analogs, could offer a practical strategy to reduce the incidence of primary CDI in vulnerable inpatient cohorts.

Beyond LXA_4_, other SPMs, including resolvins, protectins, and maresins, represent promising candidates for modulating host responses in CDI. These mediators exhibit diverse pro-resolving and tissue reparative functions that could be harnessed to prevent or treat infection-driven intestinal damage. Future studies should systematically evaluate SPM libraries in relevant preclinical models to identify compounds with optimal efficacy, pharmacokinetics, and safety profiles for clinical translation.

Looking forward, the presented GOC-based CDI model offers a robust platform for studying host–pathogen interactions, yet its physiological relevance could be further enhanced by incorporating key components of the gut ecosystem. Integration of commensal microbiota would allow investigation of microbial community dynamics in shaping CDI susceptibility and the efficacy of SPMs such as LXA_4_. Additionally, inclusion of immune cells, such as macrophages or neutrophils, would enable more comprehensive analysis of mucosal immune responses and the immunomodulatory mechanisms of SPMs. Such an advanced system would better recapitulate the intestinal microenvironment, providing a more holistic tool for dissecting the multifaceted mechanisms of CDI pathogenesis and resolution.

## Methods and Materials

### Fabrication of the GOC microfluidic device

The microfluidic chip was fabricated using soft lithography. Briefly, a template featuring raised SU-8 photoresist microstructures was first created on a silicon wafer via UV lithography. Polydimethylsiloxane (PDMS) prepolymer was prepared by mixing the base and curing agent at a 10:1 weight ratio (w/w) (Dow Corning). The thoroughly mixed prepolymer was then poured onto the template, degassed under vacuum to remove bubbles, and cured at 80 °C for 30 minutes to obtain a PDMS block containing the embossed microchannel structures. The upper and lower microchannels of the chip were designed with identical dimensions of 1 mm (width) × 10 mm (length) × 0.2 mm (height). Inlet and outlet ports were punched prior to bonding. The two channels were separated by a porous PDMS membrane. To fabricate the membrane, a silicon wafer template with an array of cylindrical pillars (10 m in diameter, 30 µm in height, 25 µm spacing) was fabricated using SU-8 photoresist. The template was then surface-treated with a silanizing agent. PDMS prepolymer was poured onto the wafer, and after degassing, a flat PDMS block was pressed onto the pillar array to allow the prepolymer to fill the inter-pillar spaces without covering the pillar tops. Curing was performed at 80 °C for 30 minutes to obtain a through-pore PDMS membrane. The upper PDMS layer was plasma-treated and bonded to the membrane; the bonded stack was then peeled off and finally sealed to the lower PDMS layer via plasma treatment ***Huh et al. (2013***).

### Cell culture and seeding in the GOC microfluidic device

Caco-2 cells were maintained in T25 flasks using Dulbecco’s Modified Eagle Medium (DMEM, Gibco) supplemented with 10% fetal bovine serum (FBS, Gibco) and 1% penicillin–streptomycin (Gibco) at 37 °C in a humidified 5% CO_2_ incubator, and passaged at a 1:3 ratio every 2–3 days. Before cell seeding, the PDMS chip was treated with ultraviolet light and ozone for 30 minutes. Microchannels were then coated with 30 µg/mL collagen type I (Sigma) for 1 hour at 37 °C. Caco-2 cells were resuspended in DMEM containing 20% FBS and 1% penicillin–streptomycin at a density of 1×10^7^ cells/mL and seeded into the upper microchannel. The chip was kept static in a CO_2_ incubator at 37 °C for 1 hour to allow cell attachment. Thereafter, cells were perfused with medium at a constant flow rate of 30 µL/h for 3 days to form a confluent epithelial monolayer.

### Culture and infection with *C. difficile* in the GOC microfluidic device

The obligate anaerobe *C. difficile* strain 1482 was routinely cultured on brain heart infusion (BHI) agar or in BHI broth under anaerobic conditions (90% N_2_, 5% CO_2_, 5% H_2_). To induce CDI on-chip, an overnight bacterial culture was diluted in fresh medium to an OD_600_ of 0.4. All equipment was placed inside an anaerobic chamber. Under static conditions, bacterial suspension (MOI = 1:10) was introduced into the upper channel (designated as 0 hours). After a 30 minutes settlement period, the chip was transferred to the transparent anaerobic chamber containing an anaerobic gas-generating sachet and an oxygen indicator (AnaeroPack™ System, Mitsubishi Gas Chemical). Tubing from medium-filled syringes was connected to the chip inlets through gas-tight holes on the side-wall of the chamber. Then the chamber was sealed by locking the top lid. The entire assembly was transferred from the anaerobic chamber to a standard cell culture incubator, where medium perfusion (30 µL/h) was resumed and maintained for 24 hours.

For LXA_4_ pretreatment experiments, the upper channel medium was replaced with anoxic BHI medium containing 200 nM LXA_4_ (Cayman) 6 hours before infection and perfused for 6 hours. For the control group, anoxic BHI medium without LXA_4_ was perfused during the same period.

For antibiotic treatment experiments, 12 hours after bacterial inoculation, the upper channel medium was switched to anoxic BHI medium containing vancomycin (0.25 or 0.5 µg/mL, Sigma) and perfused for an additional 12 hours. Control groups received anoxic BHI medium without antibiotic for the same duration.

### Quantification of bacterial load in the GOC microfluidic device

To quantify adherent *C. difficile*, the upper channel was gently flushed with 1 mL PBS at a flow rate of 50 µL/min. Effluent was collected, serially diluted, fixed, and stained with DAPI. Bacterial counts were determined using flow cytometry.

### Immunofluorescence staining

For ZO-1 staining, the upper microchannel was washed with PBS and fixed with 4% paraformaldehyde for 15 minutes, followed by PBS rinses. Cells were permeabilized with 0.25% Triton X-100 in PBS for 10 minutes, blocked with 5% BSA in PBS for 1 hour, and incubated overnight at 4 °C with Alexa Fluor 594 conjugated mouse anti-human ZO-1 monoclonal antibody (Thermo Fisher, 339194, 1:100 dilution). After washing, nuclei were counterstained with DAPI (Keygen, KGA215-10).

For mucus staining, fluorescein labeled wheat germ agglutinin (WGA-Alexa Fluor 488 conjugate, Thermo Fisher, W11261) was diluted to 25 µg/mL in culture medium and introduced into the epithelial channel. After 20–30 minutes incubation at room temperature in the dark, the channel was gently washed three times with PBS (10 minutes per wash) to remove unbound WGA. Nuclei were counterstained with DAPI. All samples were imaged using a confocal laser scanning microscope.

### Cell viability assay

Cell viability was assessed using the Cell Counting Kit 8 (CCK-8, Sigma Aldrich). Briefly, CCK-8 solution was introduced into the upper channel and incubated for 2 hours. Effluent was collected, and absorbance at 450 nm was measured using a microplate reader.

### Cytokine measurement

For all experiments involving cytokine analysis, effluent from the two outlets corresponding to the upper and lower microchannels was collected separately into microcentrifuge tubes. Collection started 6 hours before *C. difficile* infection and continued until 24 hours post infection. Collected samples were stored at –80 °C until analysis.

Levels of TNF-α and IL-8 in the effluent were measured using commercial ELISA kits (Biolegend) according to the manufacturer’s instructions. Absorbance was read on a microplate reader for concentration analysis.

For each experimental condition, the reported cytokine concentration represents the average of the values obtained from the separately collected and analyzed effluents of the upper and lower microchannels of the same chip.

### Live/dead staining

A live/dead staining kit (Invitrogen, R37601) was used. Live Green (Component A) and Dead Red (Component B) dyes were mixed 1:1, then combined with an equal volume of culture medium. The mixture was perfused into the chip and incubated for 15 minutes in the dark before imaging under a fluorescence microscope.

### Murine in vivo CDI model

Germ free female C57BL/6 mice (6–8 weeks old) were obtained from the Laboratory Animal Center of the Academy of Military Medical Sciences (Beijing, China). Mice received an antibiotic cocktail (0.4 mg/mL kanamycin, 0.035 mg/mL gentamicin, 0.035 mg/mL colistin, 0.215 mg/mL metronidazole, 0.045 mg/mL vancomycin) in drinking water for 6 days, followed by a single intraperitoneal injection of clindamycin (10 mg/kg) one day before infection. Mice were then inoculated with *C. difficile* (1×10^9^ CFU per mouse) via oral gavage. For LXA_4_ prophylaxis, mice received drinking water containing 200 nM LXA_4_ starting one day before infection. Fecal pellets were collected on the third day post infection, homogenized in PBS, serially diluted, and plated on cycloserine cefoxitin fructose agar (CCFA) for bacterial enumeration. Mice were euthanized on the third day, and intestinal tissues were collected and fixed in 10% formalin for histopathological analysis.

### Histopathological analysis

Formalin fixed intestinal tissues were embedded in paraffin, sectioned at 5 µm thickness, and stained with hematoxylin and eosin (H&E). Blinded scoring was performed to evaluate immune cell infiltration (0–3), mucosal damage (0–3), and submucosal edema (0–2). The total score (range 0–8) was classified as follows: 0–1, normal; 2–4, mild; 5–8, severe.

### RNA isolation and sequencing

Intestinal epithelial cells were harvested from the microchannels by cold PBS rinses. Total RNA was extracted using TRIzol reagent (Invitrogen) according to the manufacturer’s protocol. RNA concentration was measured with a Qubit 2.0 Fluorometer (Invitrogen, Q32866) and the Qubit RNA Assay Kit (Life Technologies, Q32855). RNA integrity and genomic DNA contamination were assessed by agarose gel electrophoresis. After quality control, RNA was quantified using the Qubit RNA Broad Range Assay Kit (Life Technologies). Strand specific mRNA libraries were constructed from 100 ng total RNA using the Hieff NGS™ MaxUp Dual Mode mRNA Library Prep Kit for Illumina^@^ (YEASEN). Libraries with insert sizes of 300–500 bp were quantified and sequenced on an Illumina HiSeq X Ten platform.

### RNA seq data analysis

Raw reads were quality trimmed using Trimmomatic (v0.36). Clean reads were aligned to the Homo sapiens reference genome (NCBI) using HISAT2 (v2.1.0). Reads mapped to exonic regions were counted with StringTie (v1.3.3b). Differentially expressed genes (DEGs) were identified using the DEP R package (limma based) with thresholds of P <= 0.05 and |log_2_(fold change)| >= 1. Gene Ontology (GO) enrichment analysis of significant DEGs was performed using the Metascape web tool.

### Statistical analysis

Data are presented as mean ± SD. Comparisons between two groups were performed using Student’s t test in GraphPad Prism 9 (GraphPad Software Inc.). P < 0.05 was considered statistically significant (* P < 0.05, ** P < 0.01, *** P < 0.001). Fluorescence intensity was quantified using ImageJ software.

## Acknowledgments

The work is supported by the Strategic Priority Research Program of the Chinese Academy of Sciences (Grant No.XDA0510200). All animal experiments were approved by the Ethics Committee of Shenzhen Institute of Advanced Technology and performed in accordance with the guidelines of the Institutional Animal Care and Use Committee of Shenzhen Institutes of Advanced Technology (SIAT-IACUC-20230817-HCS-JHZX-HSQ-A2296-01). The authors declare no competing interests.

---

**Figure 1—figure supplement 1.**
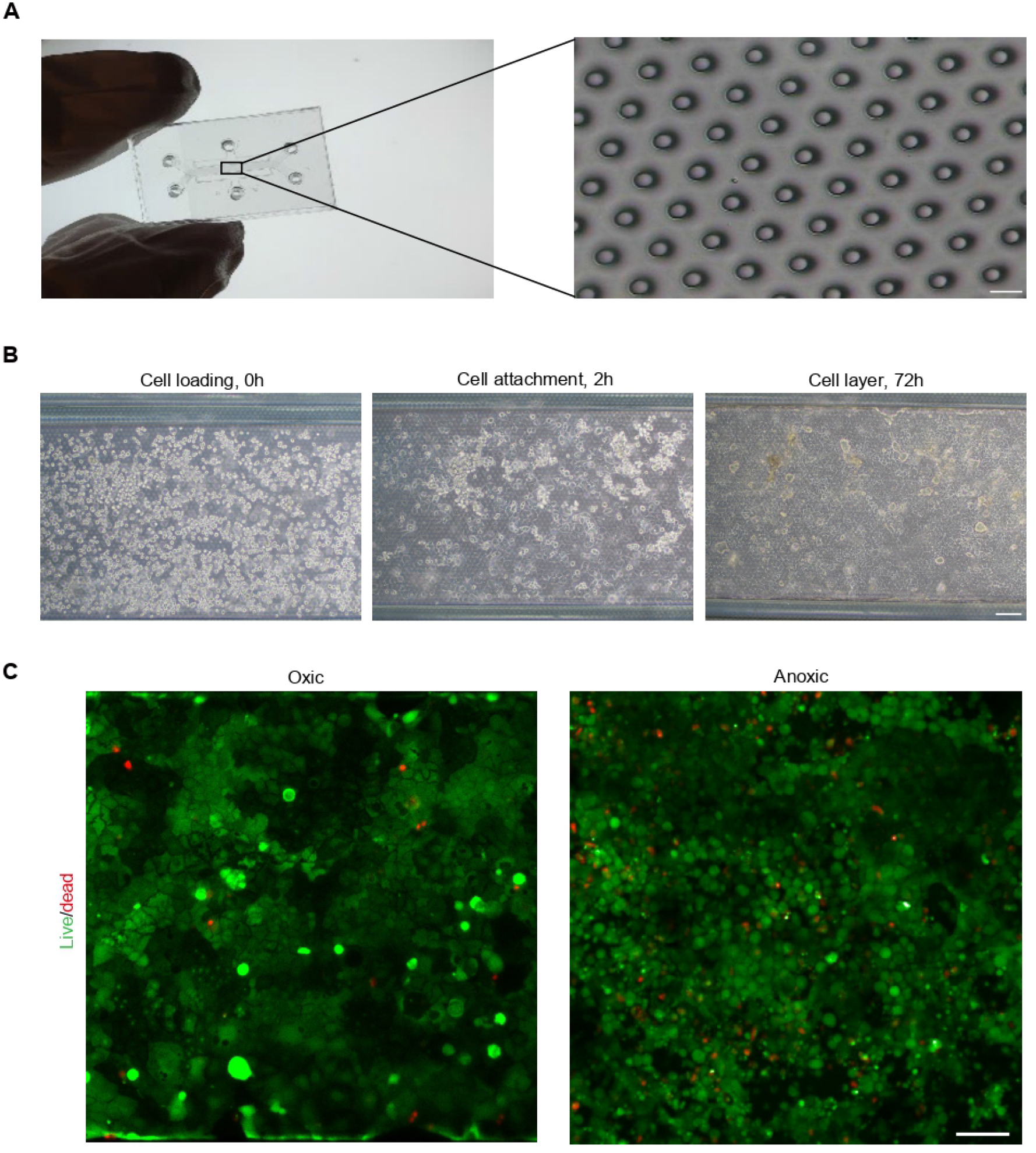
Characterization of the GOC device and on-chip cell culture. (A) Photograph of the fabricated GOC device (left) and a micrograph of its porous membrane (right). Scale bar: 20 µm. (B) Brightfield images depicted the endothelial cell layer formation procedure in the GOC. Scale bar: 100 µm. (C) Fluorescence images from a Live/Dead assay under normoxic and anaerobic conditions. Live and dead cells were stained green and red, respectively. Scale bar: 100 µm.

**Figure 2—figure supplement 1.**
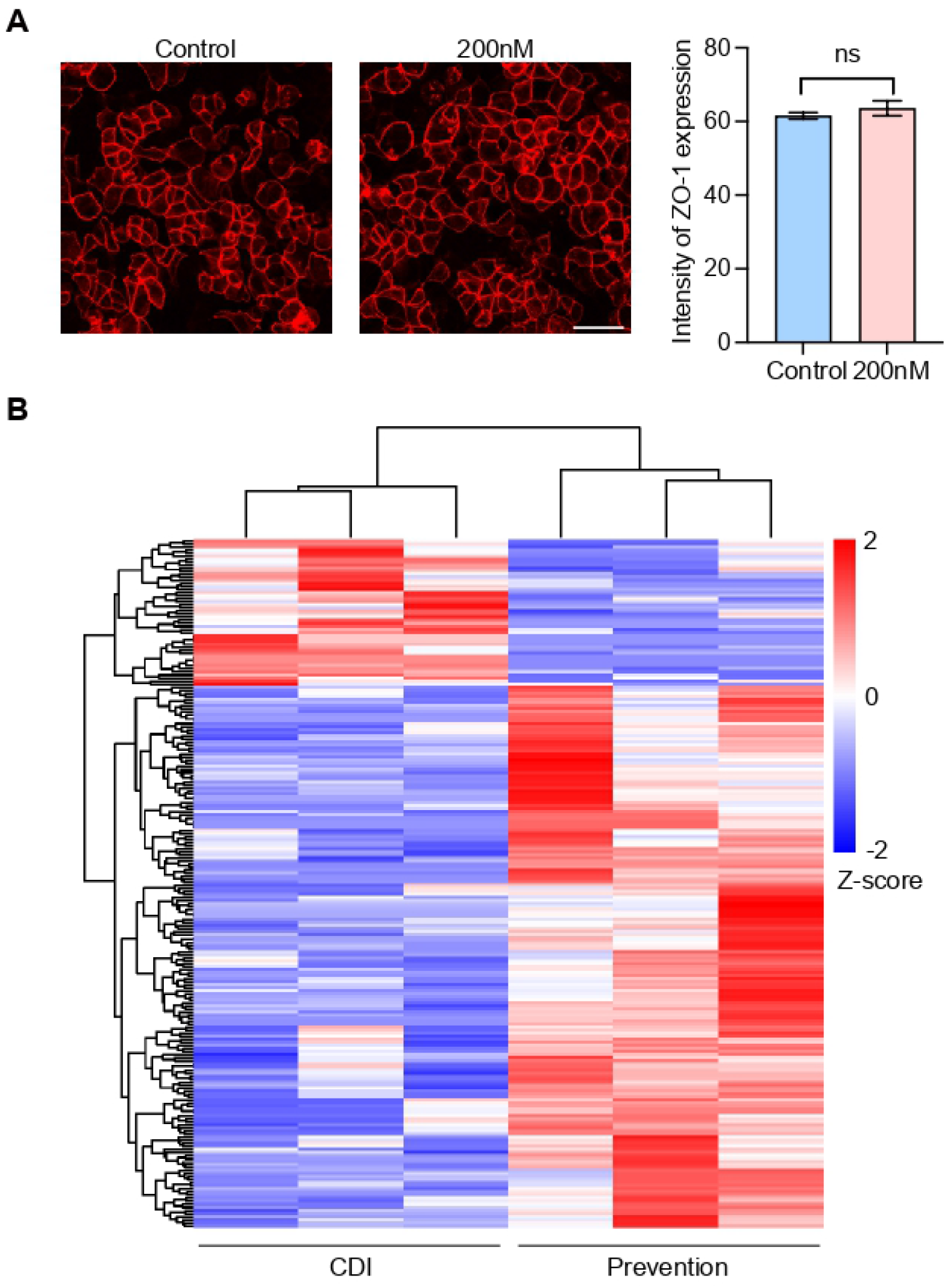
Cytotoxicity assessment of LXA_4_ and its impact on the transcriptomic landscape. (A) Confocal immunofluorescence images of ZO-1 (red) in Caco-2 cells cultured in the presence or absence of 200 nM LXA_4_. Scale bar: 50 µm. The quantitative analysis (right) shows no significant change in ZO-1 expression, indicating that LXA_4_ didn’t cause observable toxicity to epithelial cells in the GOC. Data are mean ± SD (n = 6 independent replicates). (B) Heatmap of RNA-sequencing data illustrating distinct gene expression profiles between the CDI and LXA_4_ prevention groups. Color scale corresponds to row-scaled Z-score (red: upregulated; blue: downregulated).

